# The structure and catalytic mechanism of new cellular and viral HDV ribozymes

**DOI:** 10.64898/2026.02.27.708514

**Authors:** Yuhang Luo, Xiaoxue Chen, Xiaowei Lin, Wenjian Liao, Bowen Xiao, Mengxiao Li, Zhaoji Qiu, Timothy J. Wilson, Miao Zhichao, Jia Wang, Lin Huang, David M. J. Lilley

**Author notes:** These authors should be considered joint first authors (Yuhang Luo, Xiaoxue Chen, Xiaowei Lin, Wenjian Liao, Bowen Xiao).

## Abstract

We have determined the molecular structure and investigated the catalytic mechanism of two new ribozymes of the Hepatitis delta virus family, found in the nematode *Caenorhabditis briggsae* and virus *Ackermannviridae*. Crystal structures of both conform to the double-pseudoknot architecture adopted by the viral HDV ribozyme. The *C. briggsae* ribozyme has been determined both pre- and post-cleavage. In the former both nucleotides flanking the scissile phosphate are observed, along with a metal ion, and cytosine 75 N3 bound to the O5’ leaving group. The pH dependence of cleavage rate reveals a p*K*_a_ of 6.6 and together with the inactivity of a C75U mutant provides evidence for its role as general acid. In contrast to other nucleolytic ribozymes that use catalytic metal ions, reaction rate does not depend on the p*K*_a_ of the divalent metal ion. Limited adjustment of structure of the active center is consistent with direct bonding of the metal ion to the O2’ and non-bridging O, suggesting that the ion acts as a Lewis acid to activate nucleophilic attack. This mechanism appears to be general for the HDV ribozyme class, and distinguishes it from the majority of nucleolytic ribozymes that use general base catalysis.

## INTRODUCTION

RNA catalysis is important from a number of perspectives. Some key reactions in the cell are catalyzed by RNA including mRNA splicing ^1^, and peptidyl transferase ^2,3^. Some ribozymes are widely encoded in genomes ^4-7^. According to the RNA World hypothesis ^8,9^, RNA catalysis would have played a key role in the origin of life on the planet. Yet the chemical origins of RNA catalysis are incompletely understood.

In broad terms, ribozymes catalysing phosphoryl transfer reactions divide into two major classes mechanistically. The larger ribozymes exemplified by the group I intron ribozyme ^10-15^ are metalloenzymes where two or three bound metal ions organize the active center and activate catalysis. The second group contains the nucleolytic ribozymes, a group of around 12 ribozymes that undergo site-specific cleavage by a transesterification reaction initiated by attack of a particular 2’-hydroxyl group on the adjacent phosphodiester linkage ^16,17^. In contrast to the intron ribozymes most nucleolytic ribozymes use general base-acid catalysis, employing nucleobases, the 2’-hydroxyl group or hydrated metal ions. However questions remain on how general this might be, and particularly with regard to the importance of general base catalysis ^18^.

Hepatitis delta virus is a human pathogenic virus that causes severe hepatitis. It is a satellite RNA of the hepatitis B virus that is required to enable viral replication in cells. HDV replicates by a rolling circle mechanism, generating single-stranded genomic and antigenomic circular RNA species. Each includes the sequence of a ∼85 nt closely similar nucleolytic ribozyme, the activities of which generate monomeric linear DNA species ^19,20^. The ribozymes were found to adopt secondary structures comprising five helical sections connected in a nested double pseudoknot structure. In common with other nucleolytic ribozymes, they undergo site-specific cleavage generated by attack of a specific 2’-hydroxyl group on the adjacent phosphorus atom to generate cyclic 2’3’-phosphate and 5’-hydroxyl groups flanking the cleavage site.

It was subsequently found that HDV ribozyme-like sequences occurred more widely. Szostak and co-workers ^21^ employed an in vitro selection method by which they isolated a number of self-cleaving RNA species from the human genome, one of which was clearly closely related to the HDV ribozyme. This was found in an intron of the *CPEB3* gene, and was conserved in all mammals. Using the double-pseudoknot architecture as the basis for structure-based searching subsequently led to the discovery of many more HDV-like putative ribozymes, from lamprey, lancelet, nematode, sea urchin, insects and even bacteria ^22,23^. Some were found to be active *in vivo*, but in general rates of cleavage were observed to be slow. For example, the human CPEB3 ribozyme was reported to cleave at a rate with *k*_obs_ = 0.012 min^-1^. However, cleavage could be accelerated by modification of peripheral sequences ^24^, indicating that slow cleavage was not intrinsic to the CPEB3 ribozyme structure. More recently Breaker and colleagues ^25^ used a computational pipeline involving the sequential use of BLAST sequence search followed by an analysis of secondary structure. In this work we focused on two representatives emerging from that analysis for this study (Figure 1), and verified their origins via NCBI BLAST. The first sequence is derived from a dsDNA virus belonging to the *Ackermannviridae* family, a bacteriophage known to infect *Enterobacteriaceae*. The second is from the eukaryotic nematode *Caenorhabditis briggsae*.

**Figure 1.**
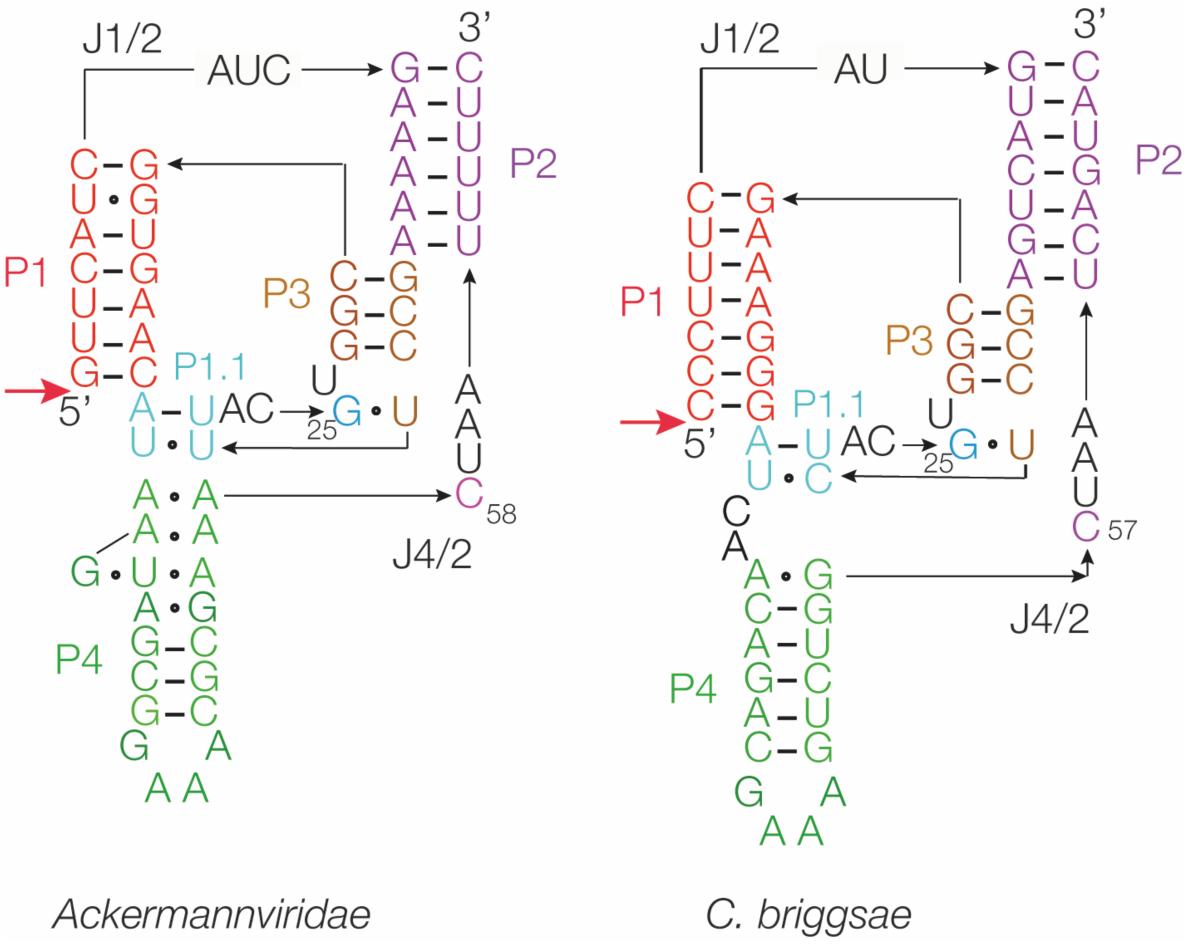
Comparison of ribozyme sequence and secondary structure for the HDV ribozyme-like sequences from *Ackermannviridae* and *C. briggsae*. The red arrows donate the predicted site of ribozyme cleavage. The expected secondary structure is colored consistently with the structural data reported here.

The structure of the viral HDV ribozyme has been determined both before and after cleavage by X-ray crystallography in a number of investigations ^26-30^. These confirmed the double-pseudoknot architecture of the RNA, with two coaxial stacks of helices (P1-P1.1-P4, and P2-P3) packed side-by-side. The catalytic mechanism of the viral ribozyme has also been studied in detail. The pH dependence of the cleavage rate indicated that a group with a p*K*_a_ = 6.6 was catalytically important ^31^, assigned to a key cytosine (C75 in the viral genomic enzyme sequence). While a C75U mutant was inactive in cleavage, activity could be restored by exogenous imidazole ^32^, and the C75 nucleobase is located adjacent to the O5’ leaving group in the crystal ^29^. This suggested a role for C75 as a general acid to protonate the 5’ oxyanion leaving group, and this was established definitively by Das and Piccirilli using phosphorothiolate substitution ^33^. The other potential participant in the catalytic chemistry is a divalent metal ion. Ions have been observed in the catalytic center of the viral ribozyme ^27,28^, and a stereospecific phosphorothioate effect was observed at the *pro*-R non-bridging oxygen atom of the scissile phosphate ^34^. In principle a metal ion might activate the O2’ nucleophile by binding as a Lewis acid, or an inner-sphere water molecule might act as a general base ^29,31,35^. This issue has not been fully resolved.

For the majority of the non-viral HDV-like ribozymes, only sequence information is available. For most there is no structural information (with the exception of some primate CPEB3 ribozymes ^36^), and there is has been almost no mechanistic investigation of any of these ribozymes. It is therefore important to explore the generality of the structural and mechanistic principles established for the viral HDV ribozyme, and we have therefore solved crystal structures for the HDV ribozymes of *C. briggsae* and *Ackermannviridae* identified by Weinberg et al ^25^. In this work we have crystallized and solved the structures of both ribozymes, both before and after the cleavage reaction in one case. We have performed detailed mechanistic investigation of these ribozymes. Our data support the role of cytosine nucleobase-mediated general acid catalysis. We observe bound Mg^2+^ ions, and the absence of a dependence of reaction rate on metal ion p*K*_a_ (in contrast to that observed for other metal ion-utilizing nucleolytic ribozymes) indicates that the major role is activation of the nucleophile as a Lewis acid.

## RESULTS

### Division of the new HDV ribozymes into ribozyme plus substrate to analyse cleavage kinetics

In this study we have investigated the three-dimensional molecular structure and catalytic mechanism of the *C. briggsae* (strain AF16, chromosome V, GenBank Accession No. CP120834.1) and *Ackermannviridae* (isolate ctUKy2, GenBank Accession No. BK037802.1) HDV-like ribozymes (Figure 1).

In the mechanistic dissection of nucleolytic ribozyme action it is experimentally advantageous to divide the RNA into two sections that can be considered to be the ribozyme and substrate, and perform cleavage reactions under single-turnover conditions. Our standard reaction conditions are 20 mM Tris (pH 7.5), 2 mM Mg^2+^ ions using 1 µM ribozyme and 150 nM substrate strands. Past experience shows that care must be exercised in how ribozymes are engineered ^37-40^, as this can readily result in a considerable loss of activity such that the catalytic chemistry may not be rate limiting. We divided *Ackermannviridae* and *C. briggsae* ribozymes according to two alternative strategies (Figure 2). These were :

1. *Breaking the connection between the P1 and P2 helices (J1/2)*. This strategy was used by Das and Piccirilli ^33^ for the analysis of the original human viral HDV ribozyme, who reported a rate of 1 min^-1^ at pH 7.5. We studied the corresponding constructs (Supplementary Table S1), generating the longer (ribozyme) strands using transcription from a PCR-generated template, and the shorter (substrate) strand by chemical synthesis. The substrate strands were radioactively [5’-^32^P]-labelled. No activity was detected for the *Ackermannviridae* HDV ribozyme. The *C. briggsae* ribozyme exhibited cleavage activity, but with an observed rate of *k*_obs_ = 0.041 ± 0.002 min^-1^ (Figure 2).
2. *Breaking the covalent continuity of helix P4*. Given the poor activity of the above forms of the new HDV ribozymes we adopted a second strategy in which the P4 helix was extended by 3 bp and the terminal loop opened (Figure 2; Supplementary Table S1). The 5’ end was extended by seven nucleotides from the expected cleavage site, so that the total length of the 5’ RNA was 56 nt, and that of the 3’ RNA was 20 nt. Both RNA species were generated by chemical synthesis. We attached a fluorophore to the 5’ terminus for detection. The labelled 56 nt strand becomes shortened to 7 nt by ribozyme cleavage. When divided in this manner the *Ackermannviridae* HDV ribozyme was now active, with an observed rate of *k*_obs_ = 4.2 ± 0.44 min^-1^ (Table 1). The *C. briggsae* HDV ribozyme exhibited strong cleavage activity, with an observed rate of *k*_obs_ = 9.3 ± 1.0 min^-1^. This is a very fast rate for any nucleolytic ribozyme, such that the chemistry is likely to be rate limiting. We have therefore concentrated our mechanistic investigation on the *C. briggsae* HDV ribozyme.

**Figure 2.**
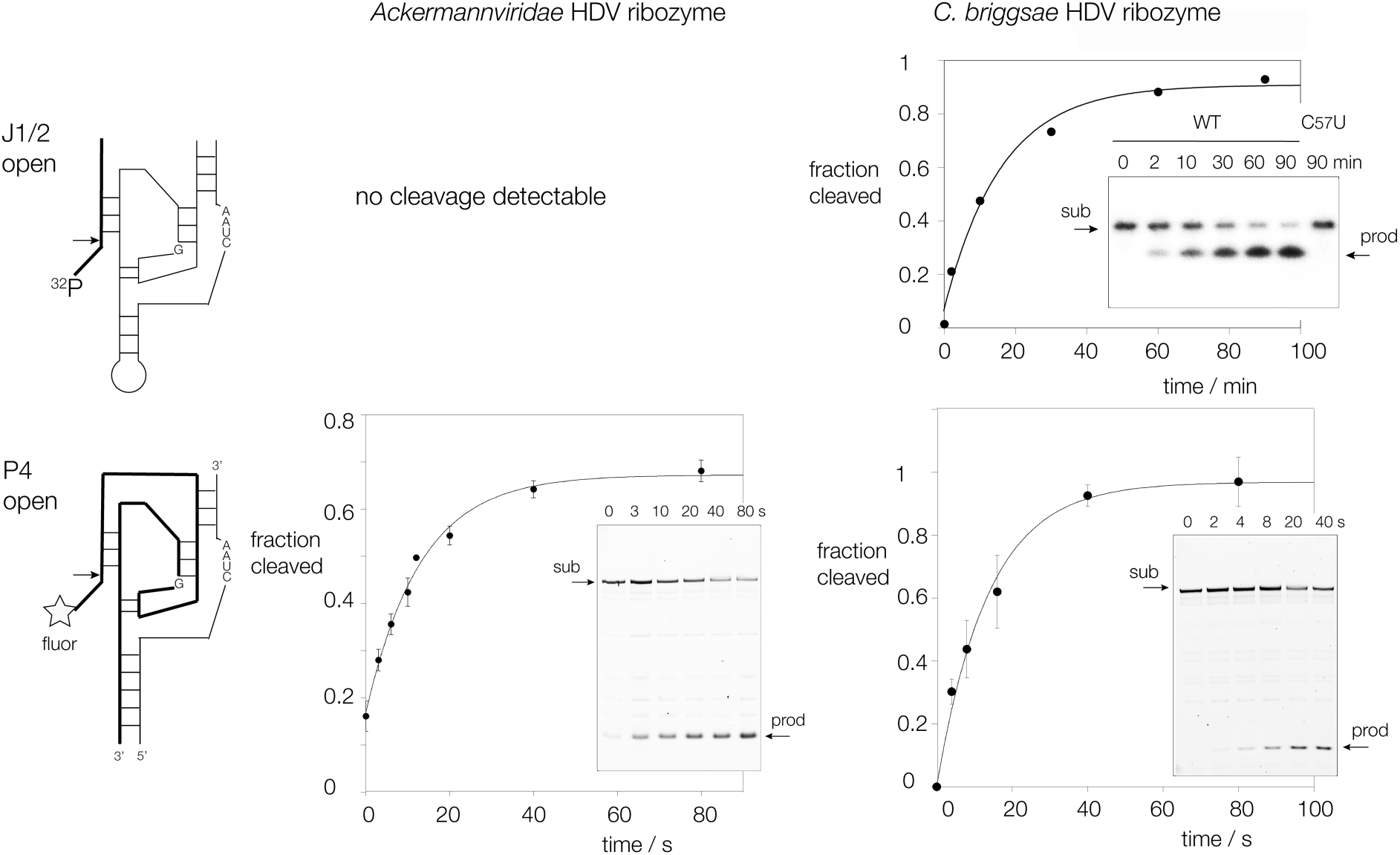
Cleavage of *Ackermannviridae* and *C. briggsae* HDV ribozymes as a function of the manner of dividing into two RNA species. In the upper constructs the RNA has been divided at the J1/2 connection between P1 and P2. In the lower constructs the P4 helix has been extended and its terminal loop opened. For each construct that exhibits cleavage activity the fraction of cleaved substrate RNA is plotted as a function of time, and the data fitted to a single exponential function. Error bars are standard deviations. The gel electrophoresis separating substrate (sub) and product (prod) is shown as an insert, with times of incubation indicated.

**Table 1.** Rates of cleavage by HDV ribozymes under single-turnover conditions. Rates were measured using ≥3 independent time courses. * other than standard conditions 20 mM Tris.HCl (pH 7.5), 2 mM MgCl_2_. N/A no activity detectable; SD standard deviation

| ribozyme | conditions * | rate<br>/ min<sup>-1</sup> | SD |
| --- | --- | --- | --- |
| *Ackermannviridae* ribozyme |  |  |  |
| J1/2 open |  | N/A |  |
| P4 open |  | 4.2 | 0.44 |
| P4 open C58U |  | N/A |  |
| *C. briggsae* ribozyme |  |  |  |
| J1/2 open WT |  | 0.044 | 0.0016 |
| P4 open WT |  | 9.3 | 1.01 |
| P4 open WT | 2 mM Ca<sup>2+</sup> | 11.1 | 1.45 |
| P4 open WT | 2 mM Mn<sup>2+</sup> | 6.7 | 1.59 |
| P4 open WT | MES (pH 6.0) | 2.4 | 0.29 |
| P4 open WT | MES (pH 6.7) | 5.1 | 0.50 |
| P4 open WT | Tris (pH 8.5) | 9.9 | 1.12 |
| P4 open C57U |  | N/A |  |
| P4 open G25O<sup>6</sup>S |  | 0.010 | 0.0016 |

### 2. Crystal structures of the new HDV ribozymes

The *C. briggsae* HDV ribozyme was crystallized as both pre-cleavage and post-cleavage structures, in hexagonal and orthorhombic space groups respectively (Supplementary Tables S2-S4). The pre-cleavage ribozyme was generated by inclusion of a 2’-deoxyribose substitution at the -1 position (so removing the potential nucleophile for the cleavage reaction), and the RNA was crystallized in the space group P6_5_22, and crystals diffracted to a resolution of 2.95 Å. The pre-cleavage ribozyme comprises five helical structures P1, P2 and P4 and two pseudoknot helices P3 and the two-base pair P1.1, all being standard A-form helices. P1, P1.1 and P4 are coaxial, as are P2 and P3, thus forming two side-by-side domains (Figure 3A, Supplementary Figure S1). These are connected by linking segments J1/2, J3/1.1, J1.1/3, J3/1 and J4/2. J4/2 includes the functionally-important C57 (corresponding to C75 in the genomic viral HDV) and the strand is located in the major groove of P3 (Figure 3B) including a single triple-base interaction between A60 and the C18:G28 base pair, making a single hydrogen bond between G28 N2 and A60 N3. Curiously, this is the only multiple-base interaction in the whole ribozyme structure. The [P1, P1.1, P4] and [P2, P3] stacked segments are also firmly connected by the formation of the P1.1 pseudoknot helix at the base of [P2, P3] (Figure 3C). The P1.1 helix comprises a standard Watson-Crick U:A base pair and a *trans* Watson-Crick C:U base pair connected by two hydrogen bonds between UN3 to CN3, and CN4 to UO4. Low rates of cleavage by some CPEB3 HDV-like ribozymes has been attributed in part to weakened stability of the P1.1 helix by inclusion of a non-Watson-Crick pairing at the second position ^21,23,24^, but the *C. briggsae* HDV ribozyme is amongst the fastest nucleolytic ribozyme despite its C:U base pair. The overall RNA structure is closely similar to that of the viral HDV ribozyme, with an RMSD = 1.77 Å (Supplementary Figure S2A).

**Figure 3.**
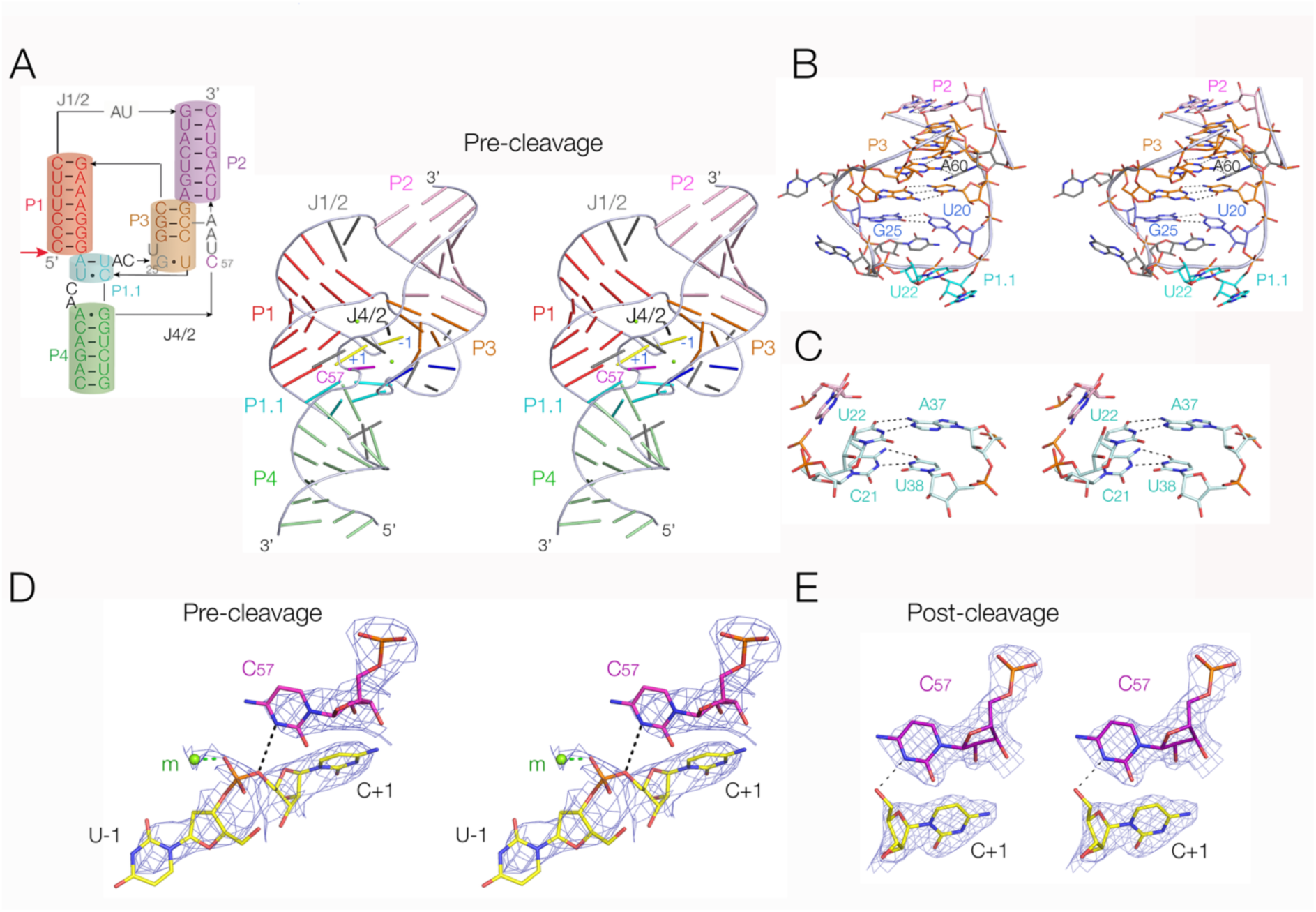
Crystal structure of the *C. briggsae* HDV ribozyme. **A.** The overall structure of the ribozyme prior to cleavage. The scheme shown on the left shows the connectivity of the helices, and is colored to match the molecular structural images. The red arrow indicates the position of the cleavage reaction. On the right the structure is shown in cartoon form with bars representing the nucleotides for clarity. The molecular graphics are shown in parallel-eye stereoscopic representation. **B.** The P2 P3 region of the structure. **C.** The P1.1 pseudoknot helix. **D.** The active site of the ribozyme prior to cleavage, with the 2**F**o-2**F**c electron density map contoured at 1.5α. Note the presence of the Mg^2+^ ion (*m*) bonded to the *pro*-R non-bridging oxygen of the scissile phosphate, and C57 N3 in position to protonate the O5’ oxyanion leaving group. **E.** The active site of the ribozyme after cleavage, with the 2**F**o-2**F**c electron density map contoured at 1.5α. A bound Mg^2+^ ion is no longer observed, but the C57 N3 remains in position adjacent to the O5’ oxygen atom.

The post-cleavage *C. briggsae* HDV ribozyme crystallized in the space group C1 2 1, and crystals diffracted to a resolution of 2.23 Å. All the structural features of the pre-cleavage ribozyme are retained in the in the structure after cleavage (Supplementary Figure S3), and the two structures can be superimposed with an RMSD = 1.27 Å (Supplementary Figure S2B).

We also crystallized the *Ackermannviridae* HDV ribozyme in its post-cleavage form. The RNA crystallized in the space group P2_1_2_1_2, and crystals diffracted to a resolution of 1.90 Å. The secondary and tertiary structure of the ribozyme (Supplementary Figure S4) is closely similar to that of the *C. briggsae* ribozyme and retains all the same conformational elements. The two post-cleavage structures can be superimposed with an RMSD = 3.41 Å (Supplementary Figure S2C). In the *Ackermannviridae* ribozyme the P1.1 helix comprises a Watson-Crick U:A base pair above a trans Watson-Crick U:U base pair connected by hydrogen bonds U21N3 to U38O2 and U38N3 to U21O4. Like the *C. briggsae* ribozyme, the non-Watson-Crick base pair does not prevent the ribozyme from cleaving at a fast rate.

### 3. The conformations of the active centers of the new HDV ribozymes

The conformations of the active center of the *C. briggsae* HDV ribozyme pre- and post cleavage are shown in Figure 3D and E respectively. In the structure of the ribozyme that has undergone cleavage, the nucleotide at the -1 position (5’ to the cleavage site) is not present. But in the pre-cleavage active center the positions of both U-1 and C1 and the connecting phosphate group are clearly defined by the electron density. This is unusual because in most structures of viral HDV ribozyme density for the -1 nucleotide is not visible, because nothing holds this nucleotide in place. However, in the *C. briggsae* HDV ribozyme structure the crystallographic symmetry closely pairs each ribozyme molecule, and the interaction includes mutual formation of a Watson-Crick-Hoogsteen base pair between U-1 of one molecule with A23 of the other (Supplementary Figure S5). In addition the U-1 nucleobases of each molecule are stacked together. This then fixes the position of nucleotide -1 in the pre-cleaved state, but is essentially an artifact of the crystal lattice. As we discuss below, this holds the C-1 and U1 in a conformation that would not be ideal for in-line nucleophilic attack.

In both pre- and post-cleavage structures the N3 of C57 is directed towards the O5’ atom. This is similar to that observed in the viral HDV ribozyme structures ^29^. The position of C57 is consistent with a role as general acid to protonate the leaving group of the cleavage reaction. A very similar position of the corresponding C58 is observed in the active center of the *Ackermannviridae* HDV ribozyme in its post-cleavage state (Supplementary Figure S4C).

In the active center of the pre-cleavage structure of the *C. briggsae* HDV ribozyme we observe electron density indicating the position of a bound metal ion 2.1 Å from the *pro*-R non-bridging oxygen atom of the scissile phosphate (Figure 3D). This is consistent with a directly-bound magnesium ion. A bound metal ion is not observed in either of the product complexes.

### 4. Cleavage rate as a function of pH

The anticipation that the new HDV ribozymes might use general acid-base catalysis leads to the prediction that the rate of cleavage will be dependent on solution pH. To be functional a general acid needs to be protonated, while a general base must be unprotonated. The observed rate of cleavage *k*_obs_ will therefore be given by

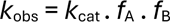

where *k*_cat_ is the intrinsic rate of cleavage (when the acid is protonated and the base unprotonated), *f*_A_ is the fraction of acid that is protonated, and *f*_B_ is the fraction of base that is unprotonated. The calculated values of *f*_A_ and *f*_B_ are plotted for the p*K*_a_ values of 6.6 (potentially a cytosine with a raised p*K*_a_ due to its environment) and 10 (i.e a high p*K*_a_ that could be a hydrated metal ion for example) respectively (Figure 4A). The profile of the cleavage reaction as a function of pH should follow that of the product *f*_A_ . *f*_B_ as shown in Figure 4B. We have measured the rate of cleavage by the *C. briggsae* ribozyme as a function of solution pH (Figure 4C). The profile is similar to that of the simulation, and the data have been fitted to this dependence. They are consistent with a lower p*K*_a_ of 6.6, and a higher p*K*_a_ > 9 that cannot be measured due to the lability of RNA at high pH. The initial rise in activity over the range pH = 6.0 - 7.5 shows that the reaction rate requires the deprotonated form of a catalytic component; this could be a general base, but would also be consistent with deprotonation of the nucleophile or solvent water for example. This increase in reaction rate at the lower pH values levels oe to form a plateau, indicating the requirement of a protonated component with an apparent p*K*_a_ of 6.6. This would be consistent with the role of C57 as general acid as suggested by its position in the crystal structures, and a closely similar pH dependence and p*K*_a_ was found for the viral HDV ribozyme by Bevilacqua and coworkers ^31^. The p*K*_a_ > 9 would be consistent with a role of a hydrated metal ion as general base, or perhaps deprotonation of solvent water that could act as a specific base. Although formally we cannot exclude the lower and higher p*K*_a_ values arising from general base and acid respectively (an example of kinetic ambiguity), this would be very much less probable in the light of the structural data.

**Figure 4.**
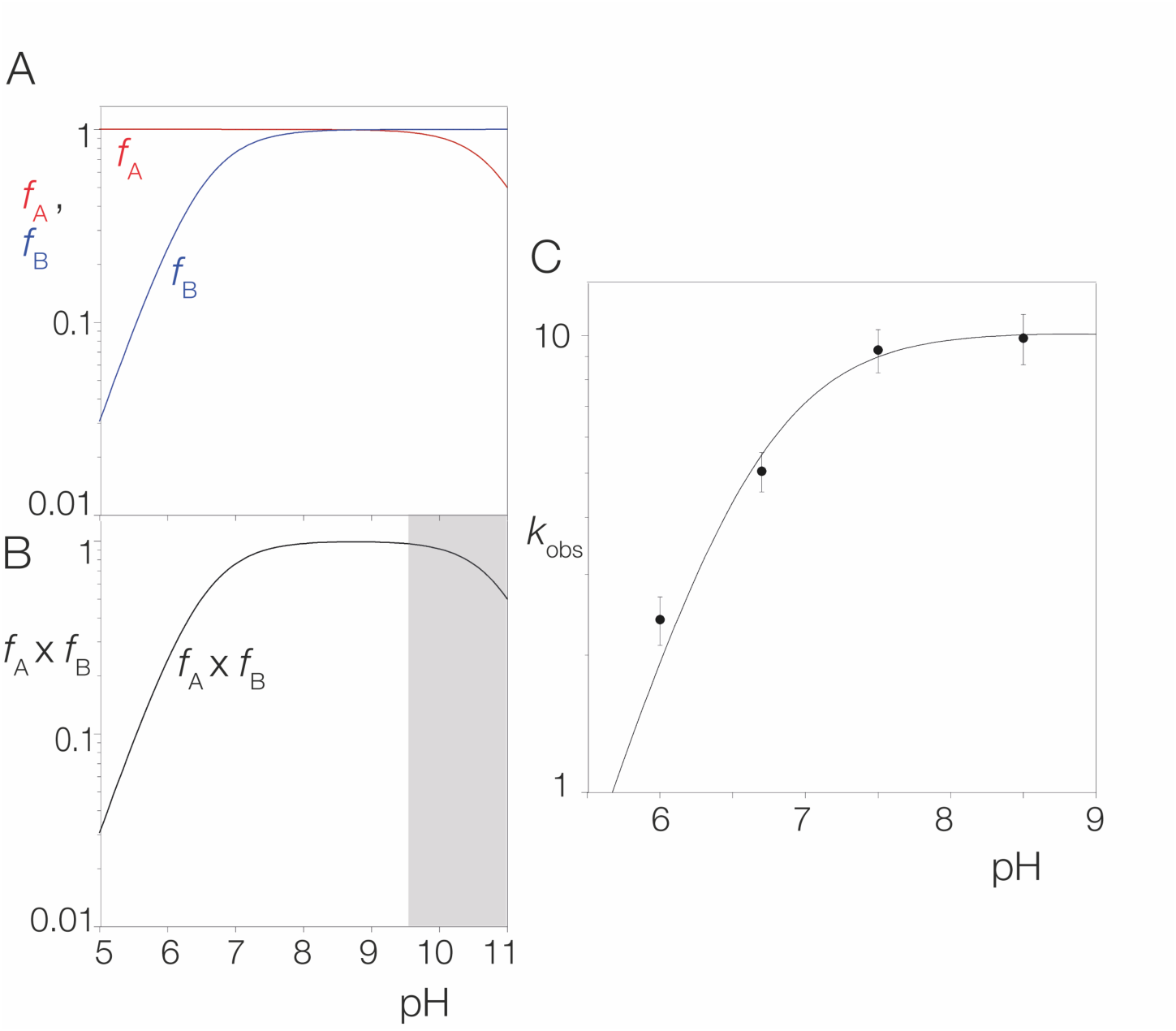
pH dependence of reaction rate. A simulation is shown on the *left*, and experimental data on the *right*. **A**. The calculated fraction of protonated acid (*f*_A_ red, calculated for p*K*_a_ = 6.6) and deprotonated base (*f*_B_ blue, calculated for p*K*_a_ = 10) plotted as a function of pH. **B**. The product *f*_A_ x *f*_B_ plotted as a function of pH. The section corresponding to pH > 9.9 is shown grey as this is not experimentally accessible due to the instability of the RNA at high pH. **C**. The experimental pH dependence of the cleavage rate for the *C. briggsae* HDV ribozyme. The measured observed cleavage rate (*k*_obs_) is plotted as a function of pH, and fitted to a double ionization process. The rates have been measured multiple times and the error bars indicate the standard deviations.

### 4. Mutation of C57, the putative general acid

The position of the nucleobase of C57 in the *C. briggsae* ribozyme (and that of C58 in the *Ackermannviridae* ribozyme) observed in the crystal structure (Figure 3D) is consistent with a role as general acid to protonate the O5’ leaving group in the cleavage reaction, as has been shown for the viral HDV ribozyme ^29^. It is also consistent with the observed p*K*_a_ of 6.6 determined above. We therefore tested this by examining the cleavage activity of a C57U mutant of the *C. briggsae* ribozyme, where the imino proton attached to C1 of the uridine is not acidic. In contrast to the wild-type sequence ribozyme, the C57U mutant exhibited no detectable ribozyme cleavage activity even after prolonged incubation (Figure 5A). No activity was also observed for the *C. briggsae* HDV C57U ribozyme divided at the J1/2 connection (Figure 2). In addition we also examined the cleavage activity of the *Ackermannviridae* ribozyme with the corresponding C58U mutation. This too exhibited undetectable levels of activity after prolonged incubation (Supplementary Fig S6). These results are consistent with the proposed role of the C57 nucleobase as general acid.

**Figure 5.**
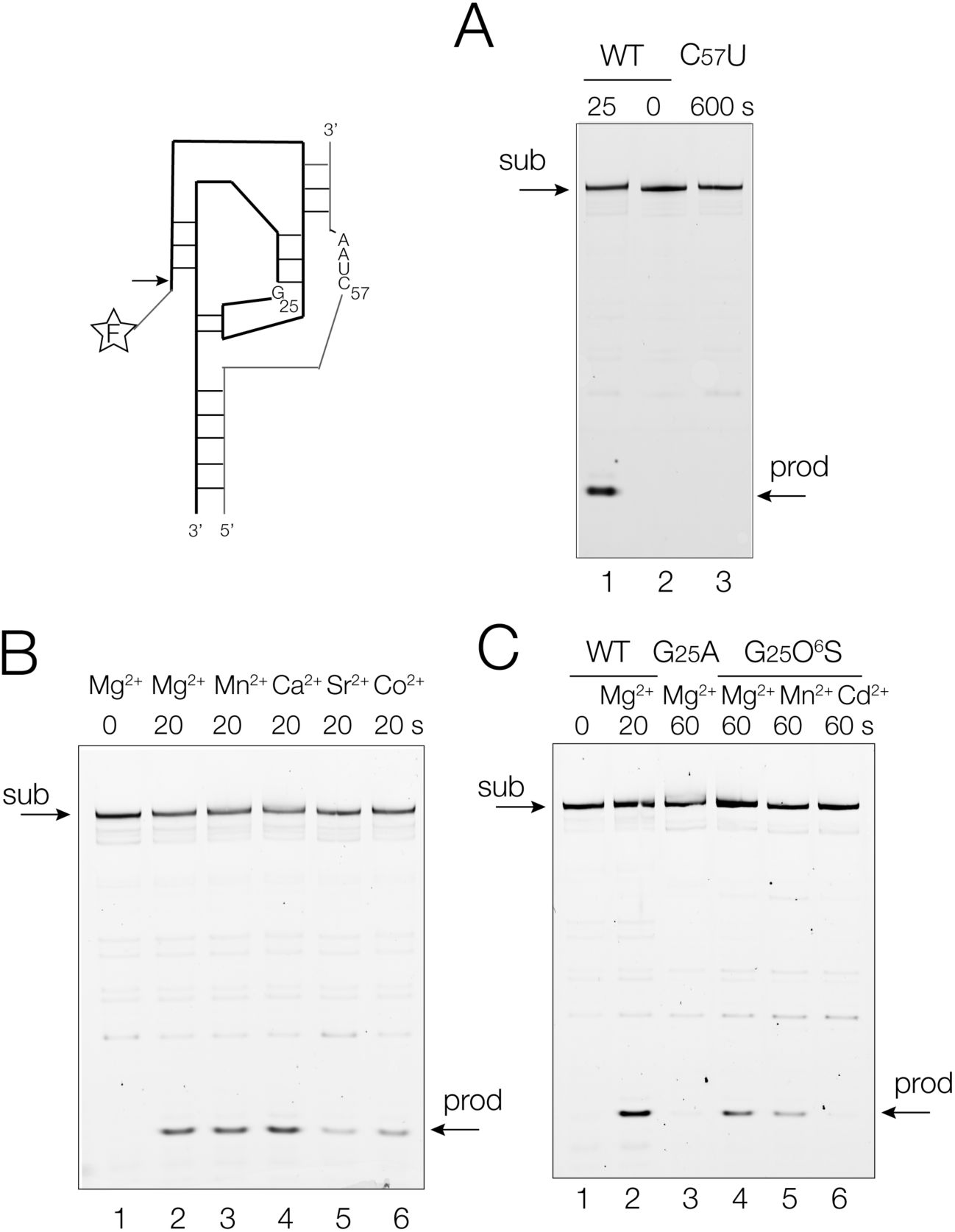
The eiect of mutation and metal ions on the cleavage activity of the *C. briggsae* HDV ribozyme. 5’-fluorescein-labelled RNA (scheme top left) was incubated under single-turnover conditions, and the substrate (*sub*) and cleaved product (*prod*) were separated by polyacrylamide gel electrophoresis under denaturing conditions. **A.** The eiect of a C57U mutation on cleavage activity. Wild-type sequence were incubated for 0 and 20 s (tracks 2 and 1 respectively). Incubation with C57U ribozyme (track 3) gave no detectable cleavage after 10 min. incubation. **B**. Activity in diierent divalent metal ions. Ribozyme and substrate were incubated under standard condition that included 2 mM of Mg^2+^, Mn^2+^, Ca^2+^, Sr^2+^ or Co^2+^ for 20 s (tracks 2-6 respectively). RNA was incubated for zero time in track 1. The rates of cleavage in Mn^2+^, Ca^2+^ (Supplementary Figure S7) and Na^+^ ions (Supplementary Figure S8) have been measured under single turnover conditions. **C**. Eiect of mutation of a putative metal ion binding site at G25. Tracks 1 and 2 respectively : incubation of wild-type RNA under standard condition that included 2 mM of Mg^2+^ for 0 and 20 s. Track 3 : incubation of G25A RNA under standard condition including 2 mM of Mg^2+^ for 60 s. Tracks 4-6 : incubation of ribozyme containing a G25O^6^S atomic mutation under standard condition including 2 mM of Mg^2+^ (track 4), Mn^2+^ (track 5) or Cd^2+^ (track 6). We have measured the rate of cleavage of G25O^6^S in the presence of 2 mM of Mg^2+^ (Supplementary Figure S9).

### 5. Activity in different metal ions

In the pre-cleavage crystal structures of the HDV ribozymes we observe a divalent metal ion bound into the active center (Figure 3D). A hydrated metal ion has been proposed as a candidate for the general base in the cleavage reaction ^31^, or it might also play a role as a Lewis acid. While the majority of the experiments described here have been performed in the presence of magnesium ions, we also investigated cleavage activity in other divalent metal ions. The *C. briggsae* HDV ribozyme was incubated in 20 mM Tris.HCl (pH 7.5) containing 2 mM Mg^2+^, Mn^2+^, Ca^2+^, Co^2+^ or Sr^2+^ ions for 20 s, and substrate and product separated by gel electrophoresis under denaturing conditions (Figure 5B). Inspection of the gel suggests that incubation in the presence of Mg^2+^, Mn^2+^ or Ca^2+^ ions leads to similar levels of cleavage activity. Lower rates of cleavage are apparent in the presence of Co^2+^ and Sr^2+^ ions. The rates of cleavage in Mn^2+^ and Ca^2+^ ions were measured (Supplementary Fig S7), and together with that in Mg^2+^ ion, the rates are all within a factor of 2, in the range of 6.7 – 11.1 min^-1^ (Table 1), in the order Ca^2+^ > Mg^2+^ > Mn^2+^. We also tested the cleavage activity of the *C. briggsae* HDV ribozyme in a high concentration of monovalent metal ion. We observed significant cleavage of the substrate RNA in the presence of 2 M Na^+^ ions, but with a rate of 0.01 min^-1^ (Supplementary Fig S8), i.e. 1000-fold slower than in the above divalent metal ions.

### 6. Mutation of G25, a potential metal ion binding site

We consistently observe a metal ion bound in the active center of the HDV ribozyme pre-cleavage structures. However, since there was no 2’-hydroxyl group at the -1 position in the crystalized ribozyme (in order to remove the nucleophile and thus prevent attack on the scissile phosphate) the position of the metal ion could be displaced from its position in the active structure. Furthermore, as discussed below, our observed structure is not in the conformation required for in-line attack, and adjustment into a geometry consistent with activity would likely lead to repositioning of the metal ion. In the viral HDV ribozyme Golden and coworkers ^28,30^ have observed a divalent metal ion bound via its inner hydration sphere to O6 and N7 of G25 that forms a G:U base pair with U20. Our observed metal ion is located >5 Å from G25 O6 and N7. We hypothesized that the metal ion might become bound to G25 in the active structure where the nucleophilic O2’ is present, and therefore examined whether or not cleavage activity might be affected by mutation at G25. We found that a G25A mutation led to a marked reduction in ribozyme cleavage (Figure 5C, track 3), with no observable cleavage after 60 s incubation. We then tested an atomic mutation in which G25 was replaced by 6-thioguanine, i.e. sulfur was substituted in place of G25 O6 (G25 O^6^S). This also led to significant impairment of ribozyme cleavage activity (Figure 5C, track 4). We measured a rate of cleavage of 0.01 ± 0.0015 min^-1^ for this variant (Supplementary Fig S9), corresponding to a 1000-fold reduction in cleavage activity.

If direct binding of a metal ion was required at G25 O6 then replacement by sulfur might weaken binding, but some activity might be restored by incubation in the presence of a softer, more thiophilic metal ion. We therefore performed further incubations of the G25 O6 ribozyme in the presence of manganese (II) or cadmium (II) ions. However, no restoration of ribozyme activity was observed on replacing 2 mM Mg^2+^ either with Mn^2+^ ions or Cd^2+^ ions (Figure 5C tracks 5 and 6 respectively).

## DISCUSSION

HDV-like ribozyme sequences have been identified in new organisms including in invertebrate species. These are very active, with rates of cleavage of 4 and 9 min^-1^, corresponding to an acceleration of cleavage of ∼ a million fold. This places the HDV ribozymes amongst the fastest of the group of nucleolytic ribozymes, comparable with the pistol and pseudoknot-containing hammerhead ribozymes ^40^. We note that the fastest ribozymes in this class are all based upon pseudoknot-containing structures, perhaps indicating that this is a particularly stable way to achieve an active fold of the RNA. In the course of this investigation we found that the HDV ribozymes can be very sensitive to the integrity of the J1/2 linker in some cases. It is therefore a safer strategy to open the P4 helix in order to convert into a two-piece ribozyme plus substrate construct.

The activity of both cellular and viral ribozymes has been retained through evolution, suggesting an important function of these ribozymes. These can now be added to the original HDV viral enzyme (for which the function in processing replication intermediates is established) and the human ribozyme CPEB1 ^21,22^ and related orthologs ^21^. Unfortunately no function has been ascribed to the cellular HDV ribozymes yet, but we feel this must exist. The sequences of these ribozymes are highly active; they have not acquired mutations disabling catalytic activity pointing to a selective advantage for preservation of activity.

Our measured pH dependence of the cleavage activity of the *C. briggsae* HDV ribozyme follows a log-linear rise in activity at lower pH that levels oe and reaches a plateau at pH 8 (Figure 4B). The data fit an apparent p*K*_a_ = 6.6, and a second significantly higher p*K*_a_. This is very similar to the pH dependence measured by Bevilacqua and colleagues for the viral HDV ribozyme ^31^. We confidently assign the lower p*K*_a_ to C57 acting as general acid to protonate the O5 oxyanion leaving group in the cleavage reaction. This is consistent with the C57U mutant exhibiting no detectable activity (Figure 5A) (and the corresponding C58U in the *Ackermannviridae* HDV ribozyme (Figure 2)), and the position of C57 or its equivalent in both ribozyme active centers shown here (Figure 3), and in those of the viral HDV ribozyme ^29^. Thus the role of the general acid in HDV ribozyme is well established, and this is clearly a general mechanism in this class of nucleolytic ribozyme. The origin of the higher (undetermined) p*K*_a_ has not been determined with confidence. In principle this might be due to deprotonation of a general base required to deprotonate the O2’ nucleophile, it might be due to deprotonation of the O2’ itself, or of solvent water. We return to this shortly.

The activity of the new HDV ribozymes depends on the presence of metal ions. Divalent metal ions (especially Mg^2+^, Mn^2+^ or Ca^2+^) lead to high rates of cleavage (Figure 5B, Table 1), but detectable cleavage activity occurred in high concentrations of Na^+^, albeit 1000-times more slowly. However, a striking result was the observation of very similar cleavage rates (within a factor of 2) for Mg^2+^, Mn^2+^ or Ca^2+^ ions. A very similar result was noted by Bevilacqua and colleagues for the viral HDV ribozyme ^31^. In the catalytic mechanism of the TS ribozyme the available evidence points to a bound Mg^2+^ ion acting as the general base to deprotonate the O2’ nucleophile. This results in a log-linear dependence of the rate on the p*K*_a_ of the water of hydration of the metal ion with unit gradient ^41^ (Supplementary Figure S10). Similarly, the pistol ribozyme exhibits a log-linear dependence of cleavage rate of metal ion p*K*_a_ with unit gradient ^42^ – in this case the evidence indicates that a bound Mg^2+^ ion acts as the general acid to protonate the O5’ oxyanion leaving group. The complete absence of a correlation of activity with p*K*_a_ for the HDV ribozymes suggests that while the metal ion is likely playing a role in catalysis, this does not involve proton transfer from its inner hydration sphere. An alternative role for the metal ion could be that it binds directly (i.e. not mediated by a water molecule) to the O2’ atom, thereby potentially increasing its nucleophilicity by acting as a Lewis acid. Golden ^30,35^ has proposed that the viral HDV ribozyme uses Mg^2+^-ion mediated Lewis acid catalysis of the cleavage reaction. Recent ^18^O kinetic isotope effect analysis of the viral HDV ribozyme indicates a dissociative transition state ^43^. This was supported by computational quantum mechanical/molecular mechanical studies that suggested a direct coordination of the metal ion with the O2’ atom. Our present results with the *C. briggsae* HDV ribozyme are in good agreement with this view of the catalytic mechanism for the new HDV-like ribozymes.

In the active center of the pre-cleavage structure of the *C. briggsae* HDV ribozyme we observe both the C-1 and U1 nucleotides, a bound Mg^2+^ ion, and the C57 nucleobase that is the likely general acid (Figure 3D). The latter is also observed in the post-cleavage structure as well as in the corresponding structure of the *Ackermannviridae* HDV ribozyme. Our pre-cleavage structure was obtained by incorporation of a 2’-deoxynucleotide at the -1 position, so removing the potential nucleophile. When we modelled the O2’ atom into our pre-cleavage structure (Figure 6A) we found that its position was significantly distant from that required for an in-line attack, although the O2’-P distance was good, at 3 Å. The local conformation is a result of a base pair formed between U-1 with an adenine in a symmetry-related molecule in the crystal lattice (Supplementary Figure S5). As a result, the measured O2’ - P - O5’ angle was 131°, i.e. 49° away from in-line attack, giving a Breaker in-line fitness parameter ^44^ of 0.58 (Figure 6B). We therefore performed a minimal manual adjustment of the local conformation, with the largest change being a 117° rotation in the torsion angle about the O3’ – C3’ bond and a smaller rotation of 39° in that about the P – O3’ bond. The conformational adjustment was straightforward with a path that encountered no intermediate steric clash. The resulting structure (Figure 6C) has an O2’ - P - O5’ angle of 172° (approximately a *g*- to *g*+ conformational change) with a corresponding Breaker in-line fitness parameter of 0.94 (Figure 6D). Thus the potentially active conformation of the ribozyme is fully accessible from our observed structure.

**Figure 6.**
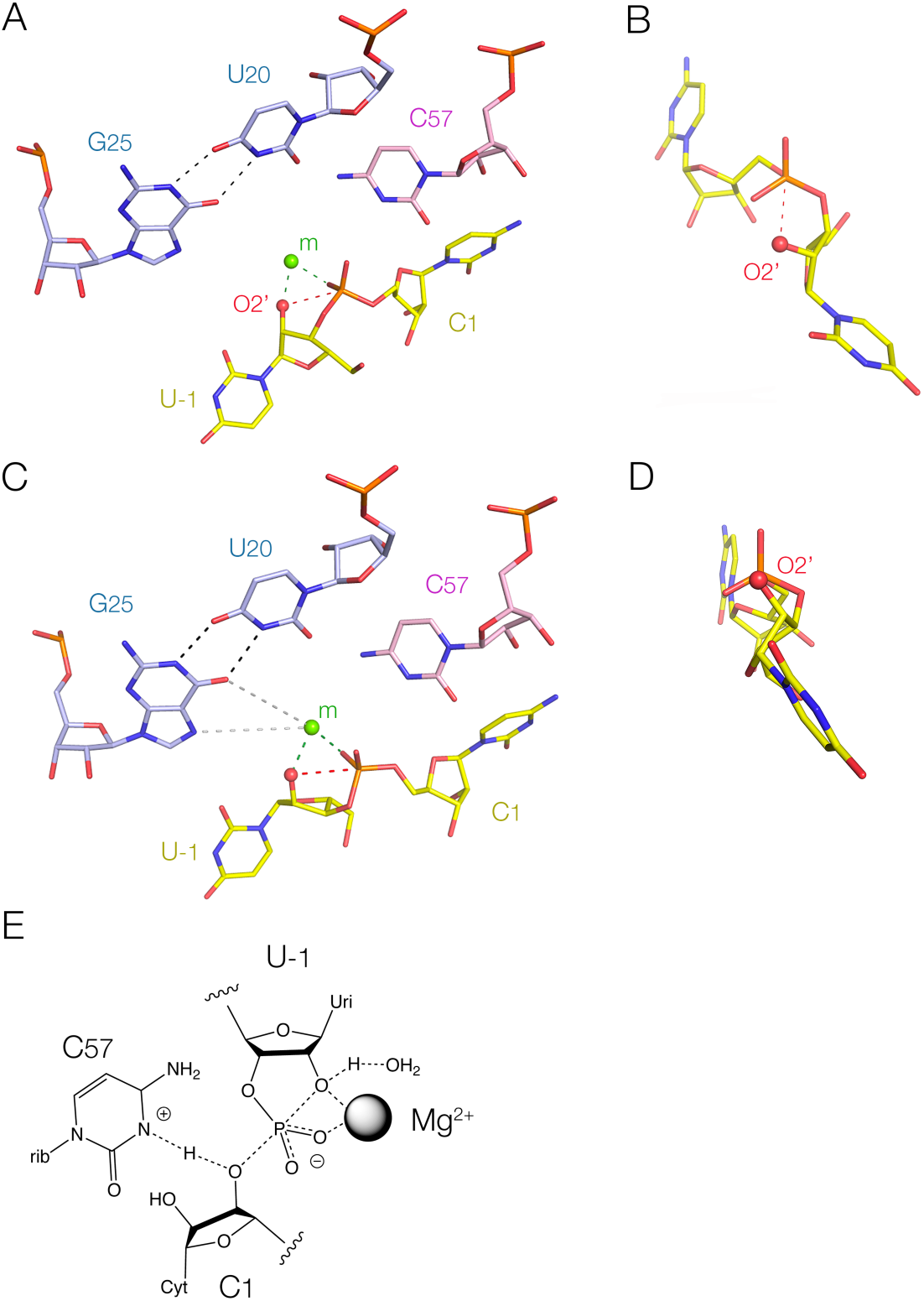
Remodelling the active center of the HDV ribozyme, and the deduced catalytic mechanism. **A**. and **B**. The active center of the *C. briggsae* HDV ribozyme with an oxygen atom modelled at the U-1 2’ position, with nothing else altered. The red broken line connects the O2’ and P atoms, representing the direction of nucleophilic attack. The bound metal ion observed in the crystal structure is shown by the green sphere. This lies 2.2 Å from the modelled O2’ atom and the *pro*-R non-bridging O atom (green broken lines). **A**. The overall view. **B**. A view down the axis of the P – O5’ bond showing the deviation from in-line geometry. **C**. and **D**. The active center of the *C. briggsae* HDV ribozyme remodelled to achieve a closer to in-line geometry, and a slightly altered metal ion position. **C**. The overall view. The metal ion lies 2.3 Å from the O2’ atom, 2.2 Å from the *pro*-R non-bridging O atom, 4.7 Å from G25 O6 and 5.2 Å from G25 N7. **D**. A view down the axis of the P – O5’ bond showing that the remodelled structure is close to an in-line geometry. **E**. The proposed catalytic mechanism of the *C. briggsae* and *Ackermannviridae* HDV ribozymes, consistent with the structural and mechanistic data. The catalytic rate enhancement arises from a combination of metal ion-mediated Lewis acid activation of the O2’ nucleophile, and general acid catalysis by C57 (C58 in the *Ackermannviridae* HDV ribozyme) protonating the O5’ oxyanion leaving group.

The Mg^2+^ ion observed in the pre-cleavage structure was directly bound to the *pro*-R non-bridging oxygen atom of the scissile phosphate at a bonding distance of 2.2 Å. This corresponds well with the stereospecific phosphorothioate effect observed with the viral HDV ribozyme ^34^. Once the O2’ atom was added to the structure, this too had a metal-O distance of 2.2 Å, consistent with direct binding to the nucleophile. However, the absence of the O2’ atom in our ribozyme as crystallized could perturb the exact position of the metal ion. In a number of structures of the viral HDV ribozyme, Golden and co-workers have observed binding of their active site Mg^2+^ ion to O6 and N7 of G25 that forms a base pair with U20 adjacent to the active center ^28,30^. In the *C. briggsae* HDV ribozyme we found that G25A mutation lead to marked impairment of cleavage activity (Figure 5C). Furthermore, the atomic mutation of G25 O6 to sulfur resulted in a 1000-fold reduction in cleavage rate (Supplementary Figure S8). That change could not be compensated by changing to a softer metal ion, either Mn^2+^ or Cd^2+^ (Figure 5C). It is therefore unlikely that the Mg^2+^ ion is directly bound to O6, and is more likely making a water-mediated interaction via its inner sphere of hydration. We therefore made a further small adjustment in the position of the metal ion in our re-modelled active center so that it is now located 4.7 Å from G25 O6 and 5.3 Å from G25 N7 (consistent with water-mediated interactions), and 2.2 Å (direct bonding) to the O2’ nucleophile and the *pro*-R non-bridging oxygen atom of the scissile phosphate (Figure 6C). This model involved very minor changes to the structure observed in the crystal, and is consistent with all the experimental mechanistic data. The data are most consistent with activation of nucleophilic attack by direct binding of the Mg^2+^ ion to both the non-bridging oxygen atom of the scissile phosphate and the O2’ nucleophile.

## CONCLUSION

The minimally-remodelled structure of the *C. briggsae* HDV ribozyme structure is consistent with all the mechanistic data, notably the reaction pH dependence, the absence of a log-linear dependence upon metal ion p*K*_a_, and mutagenesis. This indicates that the new HDV ribozymes use a catalytic mechanism involving general acid catalysis to protonate the 5’-oxyanion leaving group and divalent metal ion-mediated Lewis acid catalysis to activate attack by the O2’ nucleophile (Figure 6E). The results are also completely consistent with earlier analysis of the viral HDV ribozyme mechanism, suggesting that both structure and mechanism are well conserved in this class of ribozyme. The proposed role for metal ion-mediated Lewis acid activation of nucleophilic attack is a significant difference from other nucleolytic ribozymes that use general base catalysis. It also contrasts with those ribozymes that use hydrated metal ions as general base (TS ^41^) or general acid (pistol ^42^ and hammerhead ^42,45^). The catalytic use of a metal ion as a Lewis acid suggests that mechanistically the HDV ribozyme is intermediate in character between the nucleolytic ribozymes that use general acid-base catalysis and larger ribozymes such as the group I intron ribozymes that principally use metal ions to organize their transitions states and activate catalysis ^10-15,46^.

## MATERIAL AND METHODS

### Preparation of RNA

RNA sequences for analysis of cleavage are tabulated in Supplementary Table S1. Both strands of the *C. briggsae* and *Ackermannvirdae* HDV-like ribozymes divided at the P4 stem-loop, and the shorter (substrate) strands divided at the J1/2 connection were generated by chemical synthesis. RNA was synthesized using *t*-BDMS phosphoramidite chemistry implemented on Applied Biosystems 394DNA/RNA synthesizers as described in Wilson et al. ^47^ followed by standard deprotection and desilylation procedures ^48^. The longer (ribozyme) strands of both ribozymes divided at the J1/2 connection were generated by T7 RNA polymerase transcription from a DNA template generated by recursive PCR from synthetic DNA oligonucleotides using Q5 DNA polymerase (Master Mix, NEB). Substrate RNA in J1/2-divided RNA was radioactively [5’-^32^P]-labelled using T4 polynucleotide kinase. Substrate RNA in P4-divided RNA was 5’-fluorescently labelled with fluorescein (*Ackermannviridae*) or Cy-3 (*C. briggsae*). All RNA was purified by gel electrophoresis under denaturing conditions, and recovered by passive diffusion.

RNA for crystallisation was prepared by in vitro transcription from PCR-amplified DNA templates using T7 RNA polymerase. The resulting RNA molecules were purified by denaturing polyacrylamide gel electrophoresis (7 M urea). Full-length transcripts were visualized by ultraviolet shadowing, excised, and electroeluted using the Elutrap Electroelution System (GE Healthcare) in 0.5× Tris-borate-EDTA (TBE) buffer for 12 h at 200 V and 4°C. The RNA was then precipitated with isopropanol, washed with 75% ethanol, and resuspended in RNase-free water. Pre-cleavage HDV ribozyme RNA was purchased commercially: the short-chain section from Accurate Biotechnology (Hunan, China) Co., Ltd., and the long-chain portion from Weihuan Biotechnology (Shanghai, China) Co., Ltd., with the 5′ terminal uridine substituted with deoxyuridine (dU) to prevent self-cleavage. All DNA templates and RNA sequences used in this study are listed in Supplementary Table S2.

### Ribozyme kinetics

Standard conditions for studying ribozyme cleavage were single-turnover conditions in 20 mM Tris.HCl (pH 7.5), 2 mM MgCl_2_ at 37°C unless otherwise stated. Reactions at pH 6.0 and 6.7 were studied in 20 mM MES buffer, and reactions at pH 8.0 were studied in 20 mM Tris.HCl. pH was measured for the reac1on buffers using an Accumet AB150 pH meter (Fischer Scien1fic). Reac1ons were also studied where the divalent metal ion was replaced by 2 mM Mn^2+^, Ca^2+^, Sr^2+^, Co^2+^or Cd^2+^, or 2 M Na^+^.

150 nM labelled RNA was hybridized with 1 µM unlabelled RNA by heating to 80 °C then cooling to 25 °C at 0.2 °C per second in a thermal cycler. After preincubation at 37°C, ribozyme cleavage was initiated by addition of MgCl_2_ to a final concentration of 2 mM. Fast reactions were studied by mixing 2 µl droplets containing RNA and metal ions for 1 – 60 s, then mixing with 3 volumes of 50 mM EDTA in formamide. Slower reactions were measured in plastic tubes with aliquots were removed at chosen times and cleavage termination by addition of 3 volumes of 50 mM EDTA in formamide. Slow reactions requiring incubations > 1 h were carried out under mineral oil to prevent evaporation. Substrate and product RNA were separated by electrophoresis in 20% polyacrylamide gels under denaturing conditions, and imaged by autoradiography (^32^P) or fluorography (fluorescein or Cy3) using a Typhoon FLA 9500 fluorimager (GE Healthcare). Band intensities were quantified using ImageQuant, plotted and fitted by non-linear regression analysis to single exponential functions using KaleidaGraph (Abelbeck SoTware) to calculate rate constants.

### Simulation of pH dependence of cleavage rate

The observed rate of cleavage (*k*_obs_) will be lower than the intrinsic rate when the ribozyme is fully active (*k*_cleave_) according to :

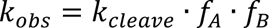

where *f*_A_ is the fraction of protonated acid, and *f*_B_ is the fraction of deprotonated base. The fractions *f*_A_ and *f*_B_ at any given pH are calculated as ^49^ :

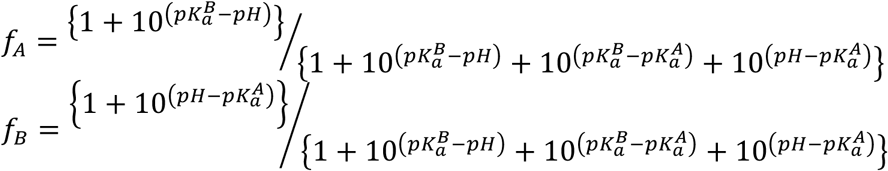

where the p*K*_a_ values of the general acid and base are written *pK_a_^A^* and *pK_a_^B^* respectively.

### Crystallisation and structure determination by X-ray crystallography

Solutions containing 10 μg/μl RNA were prepared in a buffer consisting of 5 mM HEPES at pH 7.5, 100 mM KCl, and 5 mM MgCl₂ and were denatured at 95°C for denaturation for 1 min. Crystallisation was achieved using the hanging-drop vapor diffusion method by mixing 0.8 μl RNA solution with 0.8 μl reservoir solution at 18°C. RNA sequences and crystallization conditions are summarized in Supplementary Tables S2 and S3. High-quality crystals were mounted directly on nylon loops and flash-frozen in liquid nitrogen for data collection.

X- ray diffraction data were collected at the Shanghai Synchrotron Radiation Facility (SSRF) on beamlines BL02U1 or BL10U2 and were processed using XDS ^50^. The resolution cutoff for the data was determined by examining the CC1/2 and the electron density map ^51^. The structure of the cleaved C. *briggsae* HDV ribozyme deposited as 22KS, was determined by molecular replacement using PHASER ^52^ with the previously reported HDV ribozyme structure (PDB 1DRZ) ^26^ as the search model. The resulting structure (PDB 22KS) was subsequently used as the molecular replacement template to determine the structures of the pre-cleavage C. *briggsae* HDV ribozyme (PDB 22KX) and the *Ackermannviridae* HDV ribozyme (PDB 22KW).

All models were manually adjusted using Coot ^53^ and refined through iterative cycles of refinement using Phenix.refine ^54^ and PDB-REDO ^55^. Model geometry and agreement with electron-density maps were evaluated using MolProbity ^56^ and the validation tools implemented in Coot. Simulated annealing omit maps were calculated using the composite omit map procedure in the PHENIX suite with the anneal protocol. Atomic coordinates and structure factor amplitudes have been deposited in the Protein Data Bank ^57^ under the accession codes listed in Supplementary Table S4, together with data collection and refinement statistics.

## DATA AVAILABILITY

The coordinates and structure factors of all reported crystal structures have been deposited in the PDB under the following accession numbers: 22KX (HDV-like ribozyme from *C. briggsae*, pre-cleaved state); 22KS (HDV-like ribozyme from *C. briggsae*, cleaved state); and 22KW (HDV ribozyme from *Ackermannviridae*, cleaved state).

## ACKNOWLEDGEMENTS

The authors thank the staff of the BL02U1 (https://cstr.cn/31124.02.SSRF.BL02U1) and BL10U2 (https://cstr.cn/31124.02.SSRF.BL10U2) beamlines at the Shanghai Synchrotron Radiation Facility (SSRF) for assistance in X-ray data collection.

