## Supplementary Figures and Tables for "The structure and catalytic mechanism of new cellular and viral HDV ribozymes"

#### SUPPLEMENTARY INFORMATION

#### SUPPLEMENTARY FIGURES

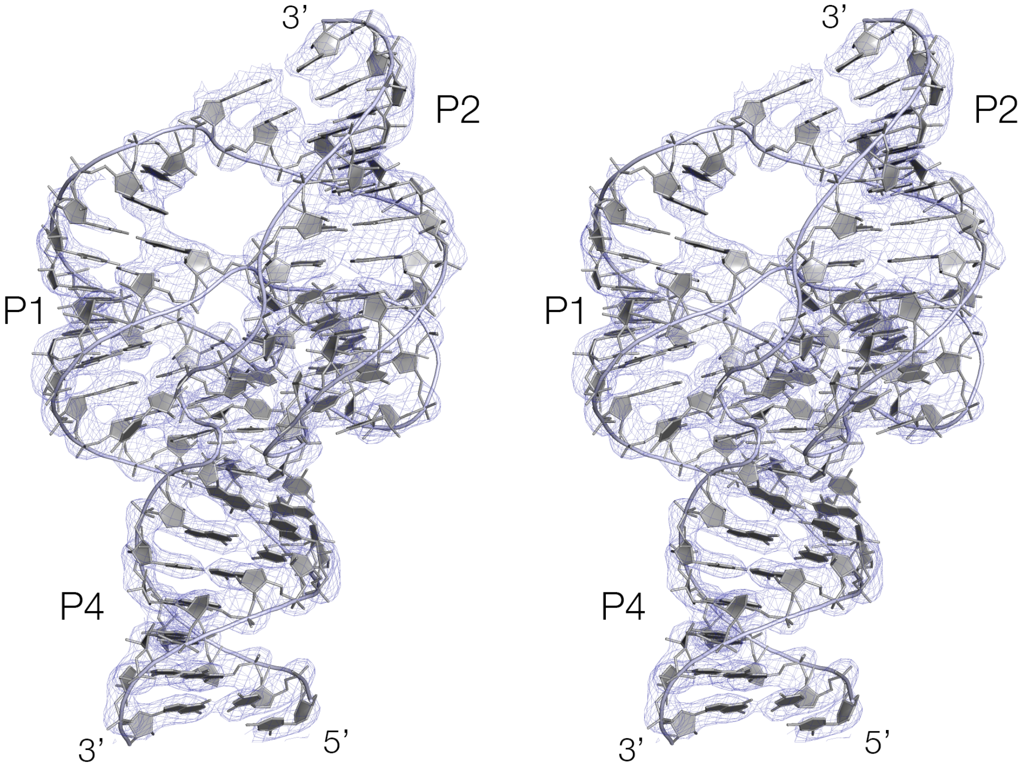


**Supplementary Figure S1**. The crystal structure of *C.briggsae* HDV ribozyme as a pre-cleavage complex showing the electron density map contoured at 1.5 σ drawn as a parallel-eye stereoscopic image.

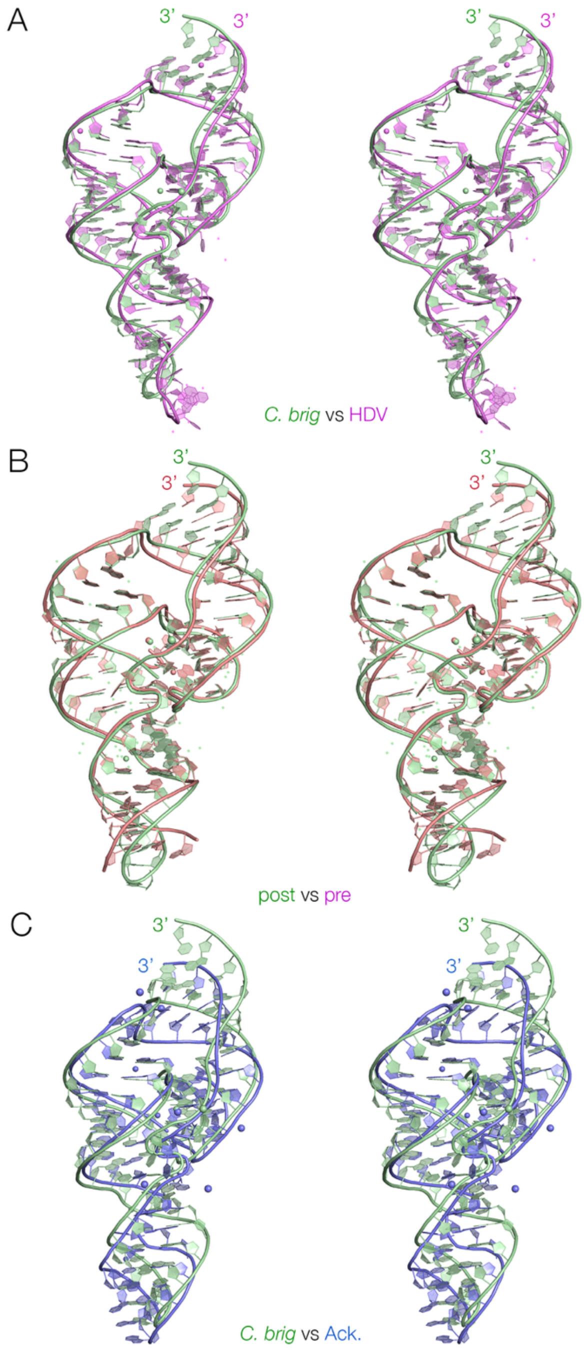


**Supplementary Figure S2**. Superposition of HDV structures.

**A**. Superposition of *C. briggsae* post-cleavage ribozyme structure (green) with the structure of the viral HDV ribozyme (violet; PDB ID 1DRZ).

**B**. Superposition of *C. briggsae* post- (green) and pre-cleavage (red) ribozyme structures.

**C**. Superposition of *C. briggsae* (green) and the *Ackermannviridae* (blue) post-cleavage ribozyme structures.

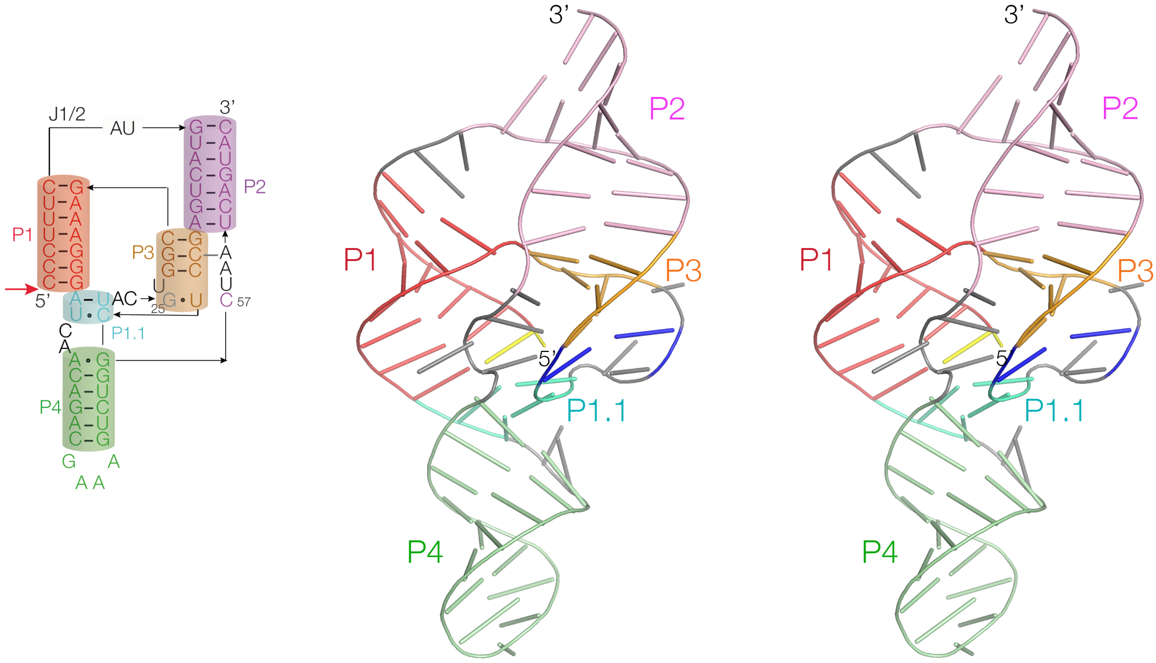


**Supplementary Figure S3**. The crystal structure of *C.briggsae* HDV ribozyme post-cleavage complex drawn as a parallel-eye stereoscopic image. The structure is depicted in cartoon mode. The 5'-terminal G1 is colored yellow and the U20:G25 basepair is colored blue. Note that the active center of the ribozyme is shown in Figure 3E.

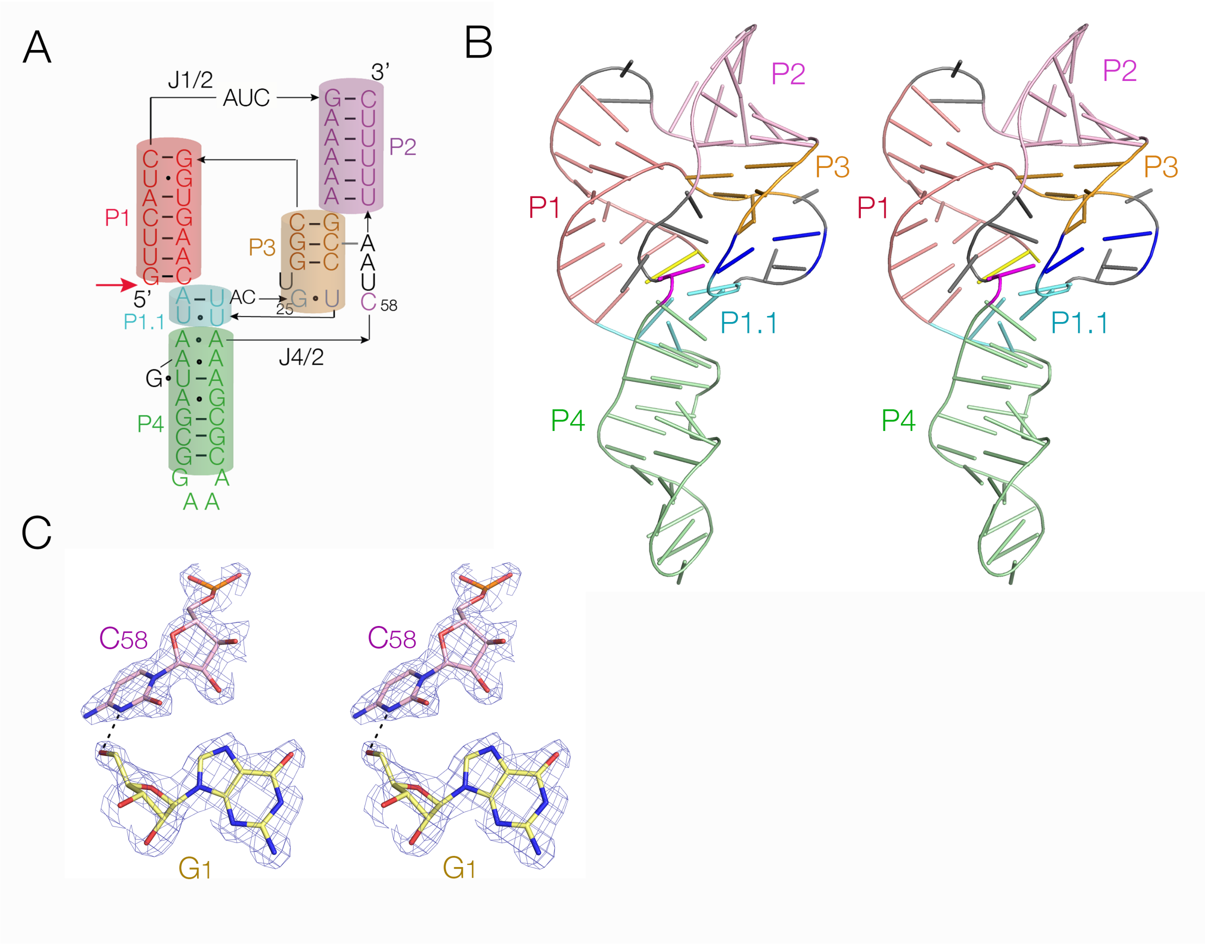


**Supplementary Figure S4**. The crystal structure of *Ackermannviridae* HDV ribozyme post-cleavage drawn as a parallel-eye stereoscopic images. **A**. Scheme of structure. **B**. The structure of the complete ribozyme depicted in cartoon mode. The 5'-terminal G1 is colored yellow, C58 is colored magenta and the U20:G25 basepair is colored blue. **C**. the active center of the ribozyme showing the electron density map contoured at 1.5 σ. Like the post-cleavage structure of the *C. briggsae* HDV ribozyme, the C58 N3 is adjacent to the O5' atom. A parallel-eye stereoscopic image is shown.

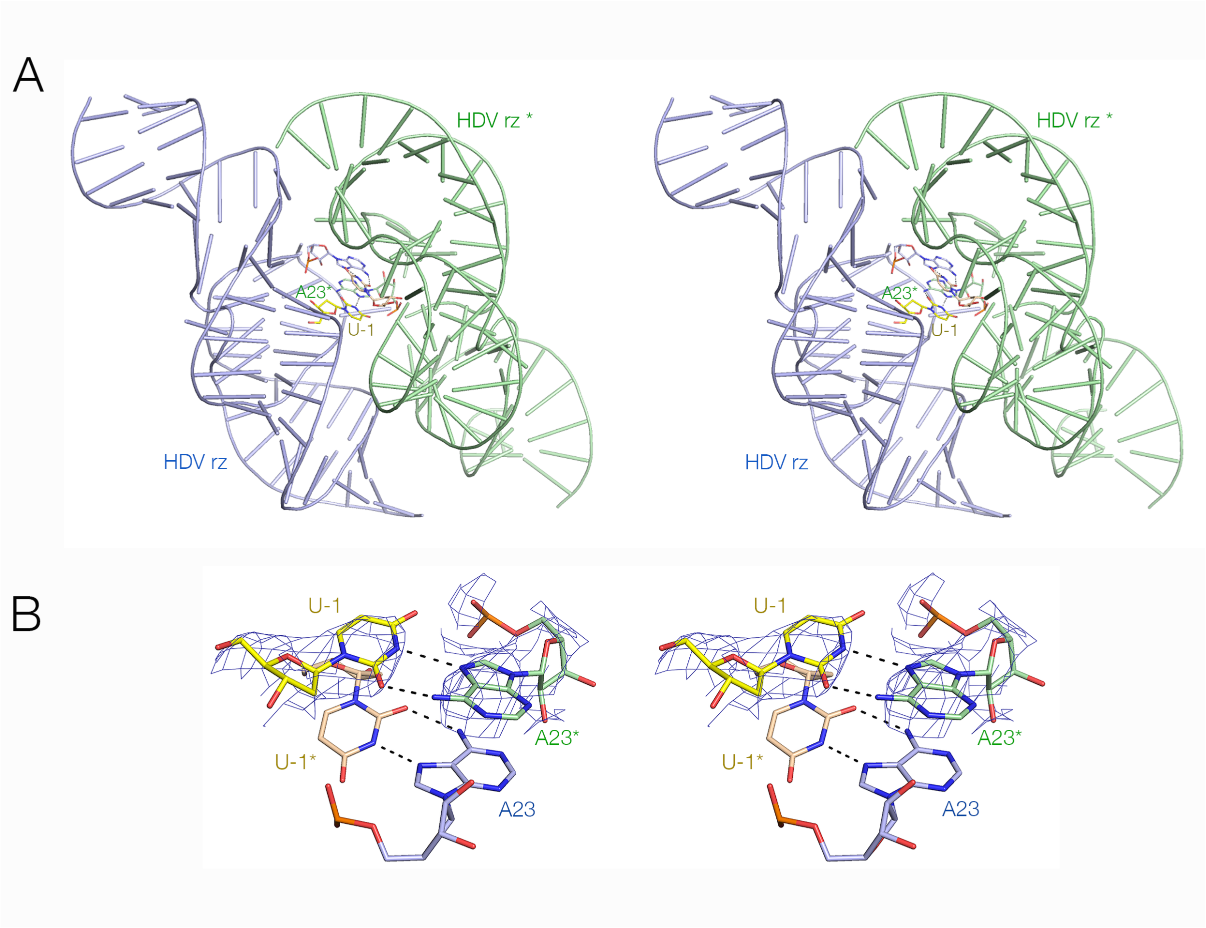


**Supplementary Figure S5**. Dimerization of the *C.briggsae* HDV ribozyme in the crystal lattice. **A**. Close interaction of two ribozyme molecules colored blue and green. **B**. Close-up view of the mutual base pairing between C-1 of one ribozyme with A23 of the other, with nucleobase base stacking between the two cytosines. The electron density contoured at 1.5 σ is shown for one of the base pairs. Parallel-eye stereoscopic images are shown.

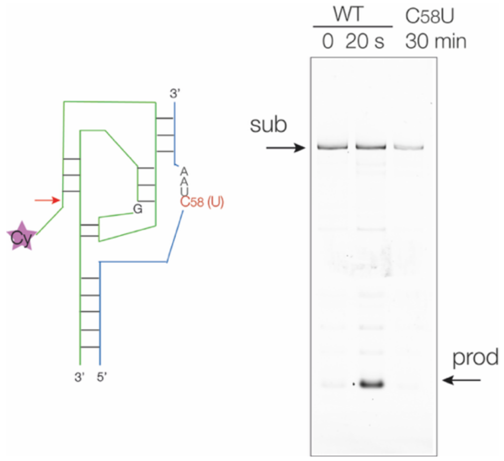


**Supplementary Figure S6**. Effect of a C58U mutation on the catalytic activity of the *Ackermannviridae* HDV ribozyme. The ribozyme was divided at the loop of the P4 helix (as shown on the left) and fluorescently-labeled with Cy3 at the 5' terminus. Incubation was performed under standard conditions containing 2 mM $Mg^{2+}$ ions at 37°C for the times shown. Note that no detectable substrate cleavage has occurred during 30 min incubation. NB in the schematic (left) the bars indicate base pairing, but not the exact number of base pairs.

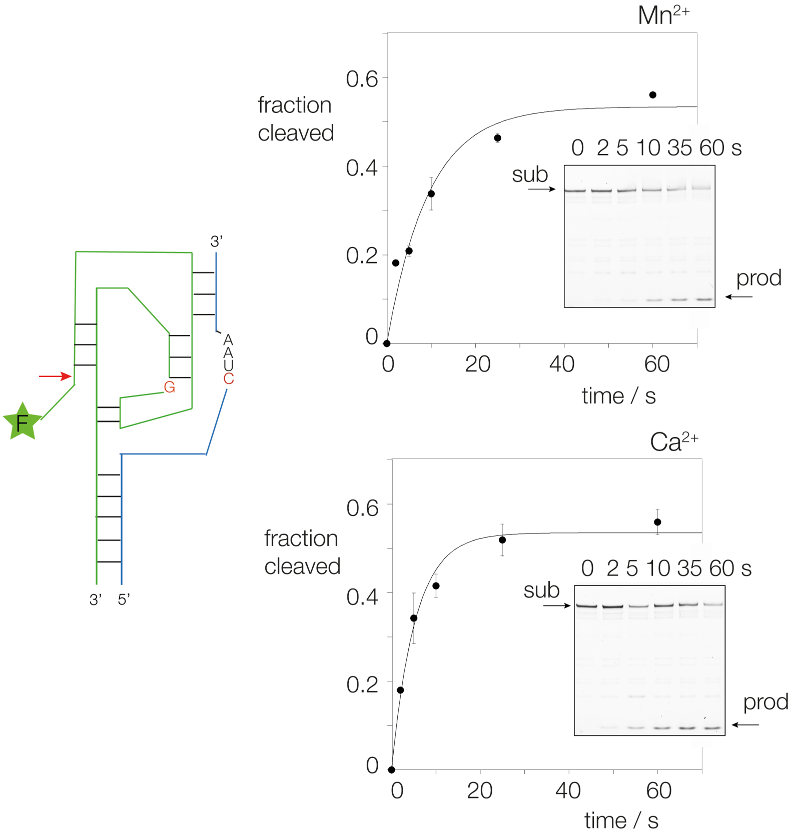


**Supplementary Figure S7**. Reaction progress of the *C.briggsae* HDV ribozyme under standard conditions containing 2 mM $Mn^{2+}$ or $Ca^{2+}$ ions at 37°C. The data have been fitted to single exponential functions giving observed cleavage rates of $k_{obs}$ = 6.7 ± 1.59 and 11.1 ± 1.45 min$^{-1}$ for $Mn^{2+}$ (upper data) or $Ca^{2+}$ (lower data) ions respectively (Table 1).

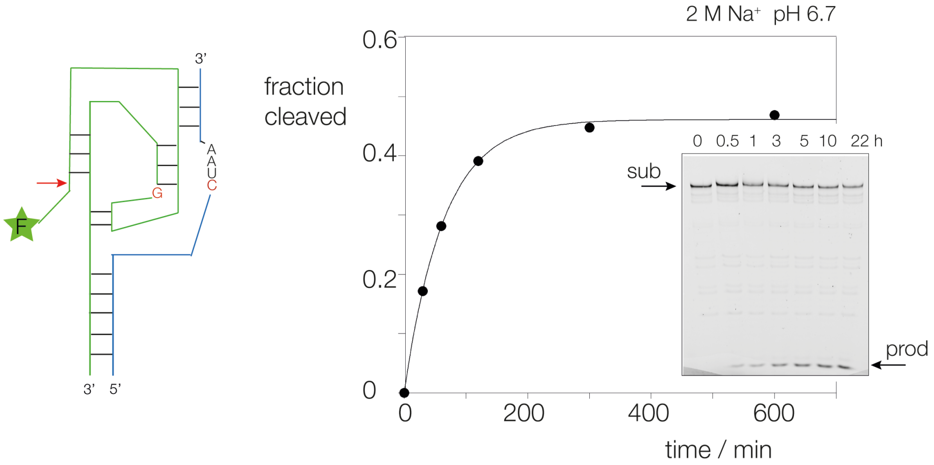


**Supplementary Figure S8**. Reaction progress of the *C.briggsae* HDV ribozyme in 20 mM MES (pH 6.7) containing 2 M $Na^+$ ions at 37°C. The data have been fitted to a single exponential function giving an observed cleavage rate of $k_{obs} = 0.01\ min^{-1}$.

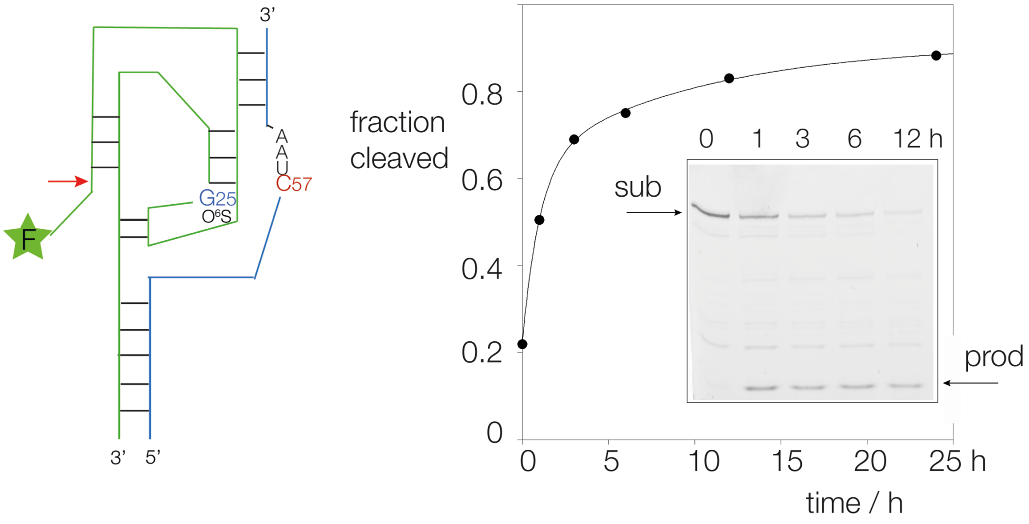


**Supplementary Figure S9**. Reaction progress for cleavage by the *C. briggsae* HDV ribozyme carrying a G25 $O^6S$ substitution under standard conditions containing 2 mM $Mg^{2+}$ ions at 37°C. The ribozyme was divided at the loop of the P4 helix (as shown on the left) and fluorescently-labeled with fluorescein at the 5' terminus. The data for this long incubation were fitted to two exponential functions, suggesting that the active conformation of the ribozyme is in slow exchange with an inactive form. The faster rate is $k_{obs} = 0.01\ h^{-1}$.

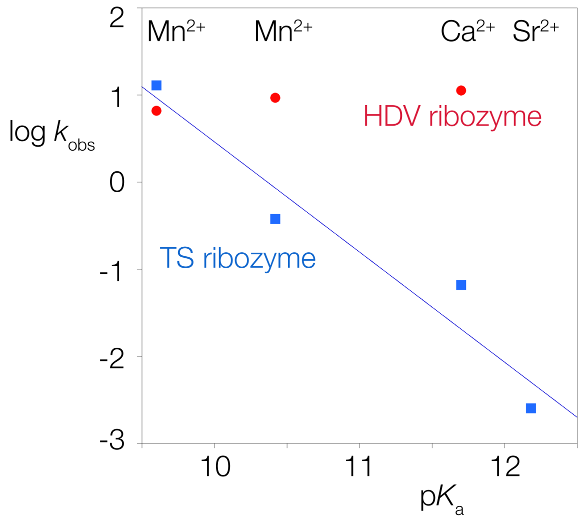


**Supplementary Figure S10**. Comparison of the dependence of observed cleavage rate vs metal ion p$K_a$ for the *C. briggsae* HDV ribozyme (red) and the TS ribozyme (blue) where the metal ion acts as general base. The data for the TS ribozyme is taken from our earlier study [Y. Liu, T. J. Wilson and D. M. J. Lilley *Nature Chem. Biol* **13**, 508–513 (2017)], and fitted to a log-linear dependence (blue line).

### SUPPLEMENTARY TABLES

**Supplementary Table S1**

J1/2 division

*Ackermannviridae*

GCAACCAGUUCAUC

GAAAAAGCCUUUACGUGGCGGUGAACAUAAGUAGCGGAAACGCGAAACUAAUUUUUC

*C. briggsae*

CCAUUUUCCCUUUC

GUACUGAGCCUCUACGUGGCGAAAGGGAUCAACAGACGAAAGUCUGGCUAAUCAGUAC

P4 division

*Ackermannviridae*

Cy3-GCAUCCAGUUCAUCAUCGAAAAAGCCUUUACGUGGCGGUGAACAUAAGUAGCGUCG

CGACGCGAAACUAAUUUUUC

*C. briggsae*

Flu-CCAUUUUCCCUUUCAUGUACUGAGCCUCUACGUGGCGAAAGGGAUCAACAGACUCGU

ACGAGUCUGGCUAAUCAGUAC

**Supplementary Table S1** Sequences of RNA molecules used in the analysis of ribozyme cleavage and kinetics. All sequences written 5' to 3'. In studying ribozyme cleavage using the J1/2 division, the short 5' strands were radioactively [5'-$^{32}$P] labelled. Using the P4 division the long 5' strands were labelled with fluorophores at the 5' termini using Cy3 (*Ackermannviridae*) or fluorescein (Flu, *C. briggsae*).

**Supplementary Table S2**

| Name | Sequence（5' seq 3'） | Notes |
|---|---|---|
| T7+6nt leader+ *C. briggsae* HDV DNA template | TAATACGACTCACTATAGCGTCGCCCCTTTCATGTACTGAGCCTCTACGTGGCG AAAGGGATCAACAGACGAAAGTCTGGCTAATCAGTAC | T7 promoter + 6-nt leader + HDV ribozyme |
| Full-length RNA transcript | GCGUCGCCCUUUCAUGUACUGAGCCUCUACGUGGCGAAAGGGAUCAACAGA CGAAAGUCUGGCUAAUCAGUAC | before self-cleavage |
| *C. briggsae* HDV ribozyme RNA | CCCUUUCAUGUACUGAGCCUCUACGUGGCGAAAGGGAUCAACAGACGAAAG UCUGGCUAAUCAGUAC | self-cleaved product |
| Pre-cleaved *C. briggsae HDV* ribozyme RNA (chemically synthesized) | NC01_dU_P4_7:（dU)CCCUUUCAUGUACUGAGCCUCUACGUGGCGAAAGGGAUC AACAGACG<br>NC01_P4-8：UCGUCUGGCUAAUCAGUAC | pre-cleaved |
| T7+7nt leader+ *Ackermannviridae* HDV DNA template | TAATACGACTCACTATAGCGTCGCGTTCATCATCGAAAAAGCCTTTACGTGGC GGTGAACATAAGTAGCGGAAACGCGAAACTAATTTTTC | T7 promoter + 7-nt leader + HDV ribozyme |
| Full-length RNA transcript | GCGUCGCGUUCAUCAUCGAAAAAGCCUUUACGUGGCGGUGAACAUAAGUAG CGGAAACGCGAAACUAAUUUUUC | before self-cleavage |
| *Ackermannviridae* HDV ribozyme RNA | GUUCAUCAUCGAAAAAGCCUUUACGUGGCGGUGAACAUAAGUAGCGGAAAC GCGAAACUAAUUUUUC | self-cleaved product |

**Supplementary Table S2.** Nucleic acid sequences used in the crystallographic study of the *Ackermannviridae* and *C. briggsae* HDV-like ribozymes.

#### Supplementary Table S3

| PDB | Crystallization conditions | cryo | nts | res | Space group | Beamline | Wavelength | Phasing |
|---|---|---|---|---|---|---|---|---|
| **HDV precleavage *Cb* NC01** | 0.2 M magnesium chloride hexahydrate,<br>0.1M HEPES sodium (pH 7.5),<br>30% v/v polyethylene glycol 400 | direct | 48+<br>19 | 2.95 | P $6_5$ 2 2 | BL10U2 | 0.9792 | MR<br>(22KS) |
| | NC01_dU_P4_7：(dU)CCCUUUCAUGUACUGAGCCUCUACGUGGCGAAAGGGAUCAACAGACG<br>NC01_P4-8：UCGUCUGGCUAAUCAGUAC | | | | | | | |
| **HDV cleaved *Cb* NC01** | 0.2 M ammonium chloride,<br>0.01 M calcium chloride<br>0.05 M Tris.HCl (pH 8.5)<br>28% w/v polyethylene glycol 4000 | direct | 68 | 2.23 | C 1 2 1 | BL02U1 | 0.9786 | MR<br>(1DRZ) |
| | CCCUUUCAUGUACUGAGCCUCUACGUGGCGAAAGGGAUCAACAGACGAAAGUCUGGCUAAUCAGUAC | | | | | | | |
| **HDV cleaved** ***Ackermannviridae*** | 0.04 M lithium chloride,<br>0.08 M strontium chloride hexahydrate,<br>0.02 M magnesium chloride hexahydrate<br>0.04 M sodium cacodylate trihydrate<br>(pH 7.0)<br>30% v/v (+/-)-2-methyl-2,4-pentanediol<br>0.012 M spermine tetrahydrochloride | direct | 67 | 1.90 | P $2_1$ $2_1$ 2 | BL10U2 | 0.9792 | MR<br>(22KS) |
| | GUUCAUCAUCGAAAAAGCCUUUACGUGGCGGUGAACAUAAGUAGCGGAAACGCGAAACUAAUUUUUC | | | | | | | |

**Supplementary Table S3.** Sequences and conditions employed in crystallization trials

*Cb : C. briggsae* ; *cryo*: cryoprotectant; *res*: resolution ; MR: molecular replacement using the structure in parenthesis.

**Supplementary Table S4**

| Name | HDV pre-cleavage *C.briggsae* NC01 | HDV post-cleavage *C.briggsae* NC01 | HDV post-cleavage *Ackermannviridae* |
|---|---|---|---|
| PDB | 22KX | 22KS | 22KW |
| **Data collection** | | | |
| Space group | P $6_5$ 2 2 | C 1 2 1 | P $2_1$ $2_1$ 2 |
| Cell dimensions | | | |
| *a*, *b*, *c* (Å) | 85.55 85.55 281.18 | 188.70 47.43 49.91 | 93.24 61.14 62.16 |
| $\alpha, \beta, \gamma$ (°) | 90.0 90.0 120.0 | 90.0 96.9 90.0 | 90.0 90.0 90.0 |
| Phasing (PDB) | MR ( 22KS ) | MR ( 1DRZ ) | MR ( 22KS ) |
| Wavelength | 0.9792 | 0.9786 | 0.9786 |
| Resolution (Å) | 140.59 - 2.95 (3.11 - 2.95) | 49.55 - 2.23 (2.35 - 2.23) | 51.72 - 1.90 (2.01 - 1.90) |
| $R_{merge}$ | 0.075 (1.766) | 0.118 (3.232) | 0.059 (0.583) |
| $R_{pim}$ | 0.018 (0.429) | 0.055 (1.589) | 0.028 (0.366) |
| $I / \sigma I$ | 20.1 (1.9) | 7.1 (0.9) | 18.1 (2.9) |
| *CC (1/2)* | 1.000 (0.820) | 0.995 (0.327) | 0.998 (0.799) |
| Completeness (%) | 92.6 (99.9) | 95.4 (89.8) | 98.9 (93.4) |
| Redundancy | 18.2 (17.7) | 5.8 (5.4) | 9.1 (5.7) |
| **Refinement** | | | |
| Resolution (Å) | 26.83 - 2.95 (3.06 - 2.95) | 31.22 - 2.24 (2.32 - 2.24) | 26.32 - 1.90 (1.97 - 1.90) |
| No. reflections | 12598 (1320) | 19943 (1489) | 28224 (2513) |
| $R_{work}$ / $R_{free}$ | 0.197 / 0.244 | 0.221 / 0.247 | 0.201 / 0.241 |
| Number of non-hydrogen atoms | 2832 | 2912 | 3077 |
| macromolecules | 2826 | 2846 | 2852 |
| ligands | 6 | 4 | 20 |
| water | 0 | 62 | 205 |
| Average *B*-factors | 130.19 | 77.89 | 50.36 |
| ligands | 118.65 | 69.84 | 39.18 |
| water | | 51.71 | 44.69 |
| R.m.s. deviations | | | |
| Bond lengths (Å) | 0.003 | 0.004 | 0.014 |
| Bond angles (°) | 0.90 | 0.93 | 1.95 |

*Values in parentheses are for highest-resolution shell.

**Supplementary Table S4.** Details of data collection and refinement statistics for the crystallographic data as deposited with the PDB.
